# MargheRita: streamlining MS-DIAL output analysis and metabolite identification in R

**DOI:** 10.1101/2024.06.20.599545

**Authors:** Ettore Mosca, Marynka Ulaszewska, Zahrasadat Alavikakhki, Edoardo Niccolò Bellini, Valeria Mannella, Gianfranco Frigerio, Denise Drago, Annapaola Andolfo

## Abstract

In the field of untargeted metabolomics, the deployment of high-resolution mass spectrometry technologies generates an immense volume of complex metabolite signals. This data density necessitates sophisticated computational frameworks for post-acquisition processing and the integration of specialized databases for accurate metabolite identification. Currently, many web-based data processing solutions offer fragmented workflows, covering only specific stages of the analysis and frequently requiring researchers to migrate data across multiple, often incompatible, platforms.

To address these challenges, we introduced margheRita, an R package designed to streamline the workflow for untargeted metabolomic profiling. Developed to work seamlessly with MS-DIAL output, margheRita provides a comprehensive pipeline for liquid chromatography-tandem mass spectrometry (LC-MS/MS) data. This tool is particularly effective for Data-Independent Acquisition (DIA) experiments, where the high-resolution acquisition of all MS/MS spectra demands rigorous and integrated processing capabilities.

A key innovation of margheRita is its ability to significantly enhance fragment matching accuracy. It achieves this by utilizing an original, curated high-quality spectral library from authentic reference standards. This library includes data acquired in both positive and negative ionization polarities using various chromatographic columns, ensuring high versatility. By bridging the gap between initial MS-DIAL processing and final biological insights, margheRita offers a holistic solution from metabolite identification to the functional interpretation of complex biological datasets.

## Introduction

Metabolomics is paving the way for comprehensive studies of the low molecular weight molecules within an organism [1]. Metabolites are regulated by both biological and environmental factors and, therefore, provide great potential to bridge the gap between genotype and phenotype. However, the exploitation of untargeted mass spectrometry (MS)-based metabolomics is slowed down by fundamental challenges in metabolite identification [1]. Despite several computational approaches exist in the field [2], validation of retention times (rt) – the time needed by a molecule to elute from a chromatographic column – and MS/MS fragmentation pattern against true reference standard is essential for confident metabolite identification [1]. This requirement is magnified in the world of untargeted studies that relies on data-independent acquisition (DIA) mode, which detects thousands of features, i.e., mass-to-charge ratio (m/z) and rt pairs. Existing computational tools cover only parts of the workflow needed to translate raw LC-MS/MS data into metabolite abundance data. This is notably hard when data is acquired through SWATH-MS technology [3]. We developed margheRita, an R package that provides crucial functionalities for the workflow of MS-based metabolomics data analysis, independently of the acquisition mode, either data-dependent acquisition (DDA) or DIA (Figure 1). This is a key advantage of the pipeline, effectively managing high-confidence identification (DDA) and comprehensive relative quantification (DIA). While DDA generates cleaner MS/MS spectra that facilitates straightforward spectral library matching, its stochastic precursor selection inherently limits the detection of low-abundance metabolites. DIA consistently captures these low-abundance compounds—which often go undetected in DDA runs—thereby enhancing metabolite coverage and virtually eliminating the missing value problem. Indeed, DIA allows researchers to improve quantitative accuracy by sampling the entire elution profile of all fragments [4]. In this way, it makes possible a retrospective analysis too, by re-analyzing old datasets years later to look for new molecules that weren’t originally of interest. To assess the performance, we applied this R package to LC-MS/MS SWATH data. Interestingly, margheRita offers the possibility to perform metabolite identification, since it is supported by an original spectral library of true reference standards (Figure 1) for given chromatographic methods. The pipeline includes functions to reliably assign metabolite ID with 3 different evidence levels, by evaluating the rt on a specific chromatographic column, the accurate m/z of both the precursor and the fragments, and the relative intensity of the fragments. In addition to various statistical and biological analyses, the pipeline supports data transfer to external packages for further metabolomics analysis.

**Figure 1:**
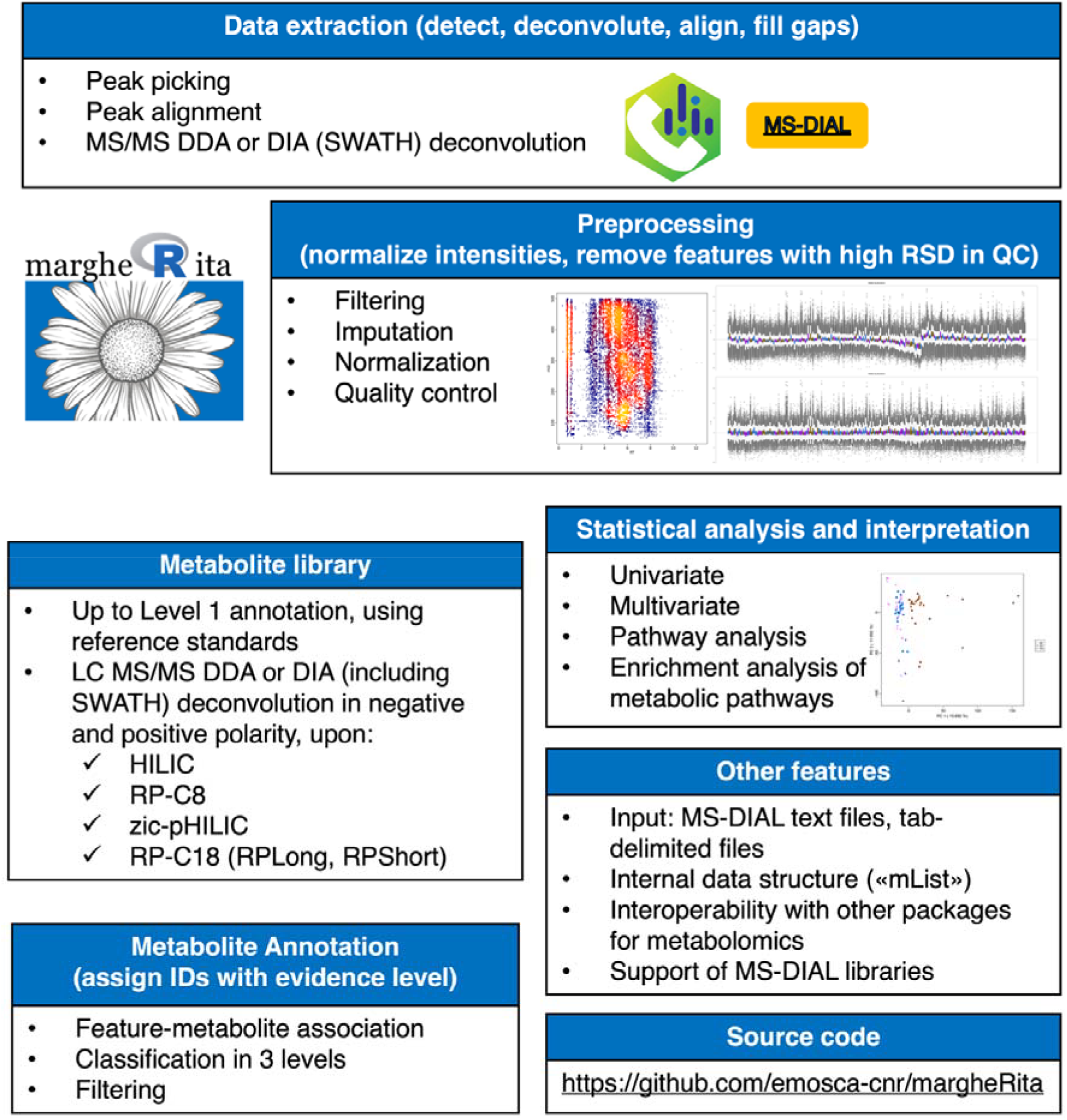
Overview of margheRita. The main functionalities of margheRita R package are summarized in each block: margheRita is dependent on MS-DIAL text files (tab delimited), which are its input files. Subsequent preprocessing of data in margheRita includes filtering, imputation, normalization and quality control. Metabolite identification at level 1 is performed thanks to a spectral library of authentic reference standards, acquired in data-dependent mode (referred as information-dependent acquisition (IDA) in SCIEX software). Each reference standard was run on four different chromatographic columns, in both polarities. Additionally, publicly available libraries, in msp format, can be used for metabolite identification in margheRita package. Regardless of the library used (in-house or publicly available), metabolite annotation level is reported in the output file. Several statistical and biological analyses are offered within the pipeline, along with the opportunity to transfer data to other packages for metabolomics analysis.

### Design and implementation

The software margheRita is implemented as an R package [5]. It requires feature-level data (rt, m/z, MS/MS spectra and abundance across samples) and sample-level data in text format. Feature-level data files are obtained from raw files upon the first step of processing through the freely available software MS-DIAL [6], which performs data extraction, peak picking, peak alignment, SWATH-MS spectral deconvolution (if required) and peak identification, supporting multiple technologies and vendors. Sample-level data files must contain a series of annotations related to the experimental design, like injection order, experimental batch and condition.

Data are structured as “mRList” objects, a specific object of type list that is the shared input/output of all functions that operate on data. To support interoperability with other packages with a focus on metabolomics, mRList objects can be reorganized as a “MetaboSet” object, which is used by “notame” [7], or as a “PomaSummarizedExperiment” object, which is used by “POMA” [8].

Upon the data import from MS-DIAL, the margheRita package performs quality control, filtering, imputation and normalization. These functions provide solutions that were devised for metabolomics data analysis as well as tools that were originally developed in other domains. Examples of functions that are tightly connected with the characteristics of metabolomics data are the “heatscatter” visualization of m/z vs rt values, the filtering of features by m/z values or by coefficient of variation (in samples relatively to quality controls), and the probabilistic quotient normalization [9]. RLE (relative log expression) plots [10], a tool initially devised for gene expression data, are included in margheRita as RLA (relative log abundance) to assess whether the normalization procedure, aimed at removing unwanted variation, has been successful.

The package provides an original metabolite library, built on authentic reference standards, that supports validated annotation (“Level 1”, see below). It was acquired in both positive and negative polarity, for 4 chromatographic columns, namely BEH Amide hydrophilic interaction liquid chromatography (HILIC), Reverse-Phase (RP)-C18 with two different gradients (RPLong and RPShort), zic-pHILIC (pZIC), RP-C8 (LipC8) (Supplementary Methods, Additional File 1) and contains about 2,000 MS/MS spectra obtained through data dependent acquisition. Moreover, margheRita supports the use of MS-DIAL reference libraries, which enable putative identification (“Level 2” or “Level 3”), for large number of molecules (∼10^^4^). Differently from previously published confidence levels [11, 12], the classification proposed in this work enables us to convey the annotation confidence with simple and standardized criteria considering the matching with rt, m/z, and MS/MS, as described below.

The association between a feature *f*_*i*_ and a library metabolite *m*_*j*_, characterized by a sequence of MS/MS peaks, respectively 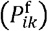 and 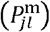 , is based on the following quantities:

i. rt error *ε*_rt_ (*f*_*i*_ , *m*_*j*_) = |*t*(*f*_*i*_) − *t*(*m*_*j*_)| , where *t* denotes rt;
ii. ppm (part per million) error 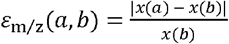 · 10^6^, where *x* denotes the is calculated on the m/z; *ε*_m/z_ feature-metabolite pair (*f*_*i*_ , *m*_*j*_) and on any feature-metabolite pair of MS/MS peaks 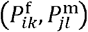;
iii. relative intensity error: 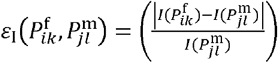 , where *I* denotes the relative intensity.

A match between *f*_*i*_ and *m*_*j*_ exists when the errors lie below the corresponding thresholds *(α*_*t*_ , *α*_m/z_, *α*_*I*_) and is classified as (Table 1):

**Table 1.** Classification levels and type of supporting evidence. The feature-metabolite match is called based on the quantity reported between brackets.

| Classification | rt<br>($\varepsilon_t$) | m/z<br>($\varepsilon_{m/z}$) | MS/MS<br>($\varepsilon_l, \varepsilon_{m/z}$) |
| --- | --- | --- | --- |
| Level 1 | √ | √ | √ |
| Level 2 |  | √ | √ |
| Level 3a | √ | √ |  |
| Level 3b |  | √ |  |

i. “Level 1” (supported by rt, m/z and MS/MS as compared to the analyzed reference standard), if *ε*_rt_ (*f*_*i*_ , *m*_*j*_) < *α*_rt_ , *ε*_m/z_ (*f*_*i*_ , *m*_*j*_) < *α*_m/z_ , and there exists a pair of feature-metabolite MS/MS peaks 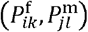 such that 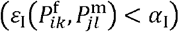 and 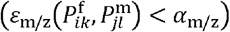;
ii. “Level 2” (supported by m/z and MS/MS), if *ε*_m/z_ (*f*_*i*_ , *m*_*j*_) < *α*_m/z_ and there exists a pair of feature-metabolite MS/MS peaks 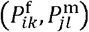 such 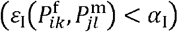 and 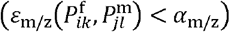;
iii. “Level 3a” (supported by rt and m/z), if *ε*_rt_ (*f*_*i*_ , *m*_*j*_) < *α*_rt_; and *ε*_m/z_ (*f*_*i*_ , *m*_*j*_) < *α*_m/z_;
iv. “Level 3b” (supported by m/z), if *ε*_m/z_ (*f*_*i*_ , *m*_*j*_) < *α*_m/z_.

In the metabolite annotation results, besides classification level, every match between *f*_*i*_ and *m*_*j*_ is provided with a number of quantitative and categorical scores (Table 2) that summarize various aspects of the supporting evidence: classification of *ε*_rt_ in {“super”, “acceptable”, “unacceptable”}; classification of *ε*_m/z_ in {“super”, “acceptable”, “suffer”, “unacceptable”}; number *n*_*ij*_ of matching MS/MS peaks; the ratio *n*_*ij*_ / *n*_*j*_, where *n*_*j*_ is the number of metabolite MS/MS peaks; whether the whole feature *f*_*i*_ is part of its set of MS/MS fragments or not. These scores are used to filter the list of associations between features and metabolites. Indeed, a series of one-to-many and many-to-one associations between *f*_*i*_ and *m*_*j*_ could arise, especially when considering large sets of features and reference metabolites. Therefore, we designed an algorithm that, leveraging the classification (Level 1, Level 2, Level 3a and Level 3b), the errors ( *ε*_rt_, *ε*_m/z_ *ε*_I_ ) and quantitative and categorical scores, resolves the vast majority of the one-to-many and many-to-one associations, retaining only those supported by the strongest evidence (Supplementary Methods, Additional File 1), thus providing a very clear, immediate and selective annotation of the features. This score system is accompanied by the plotting function to assess the similarity between the observed and reference spectra to help manual curation of data. As an example, we report the heatmaps of *ε*_m/z_ and *ε*_*I*_ values for all-pairs of MS/MS fragments between feature F3277 and Hypoxanthine MS/MS spectra (Figure 2a-b). This Level 1 identification is supported by four matches between MS/MS fragments of F3277 and Hypoxanthine (Figure 2c), *ε*_m/z_ = 4.31 (“super”, between precursor F3277 and Hypoxanthine) and *ε*_rt_ = 0.211 (“super”). It is noteworthy that the library was acquired in IDA (information dependent acquisition, which is the DDA mode on SCIEX instrumentation and in SCIEX software) mode, while the sample was acquired in SWATH-MS mode, thus explaining the differences in number and intensity of MS/MS fragments observed in the two compared MS/MS spectra (10 fragments in the sample and 9 in the library MS/MS spectrum, respectively). Even if in the SWATH experiment more precursors were contemporarily fragmented, the margheRita pipeline was able to identify the feature F3277 as Hypoxanthine.

**Table 2.** Criteria used in the feature-metabolite association process.

| Criterion | Type | Values |
| --- | --- | --- |
| Level | categorical | {1, 2, 3} |
| Level 3 note | categorical | {m/z & rt; m/z} |
| rt class | categorical | {super, acceptable, unacceptable} |
| m/z class | categorical | {super, acceptable, suffer, unacceptable} |
| rt error ($\varepsilon_{rt}$) | quantitative | $\geq 0$ |
| ppm error ($\varepsilon_{m/z}$) | quantitative | $\geq 0$ |
| Relative intensity error ($\varepsilon_l$) | quantitative | $\geq 0$ |
| Number of MS/MS matching fragments ($n_{ij}$) | quantitative | $\geq 0$ |
| MS/MS matching fragments ratio ($n_{ij}/n_j$) | quantitative | $\geq 0$ |
| Precursor included in its MS/MS spectra | categorical | {yes, no} |

**Figure 2.**
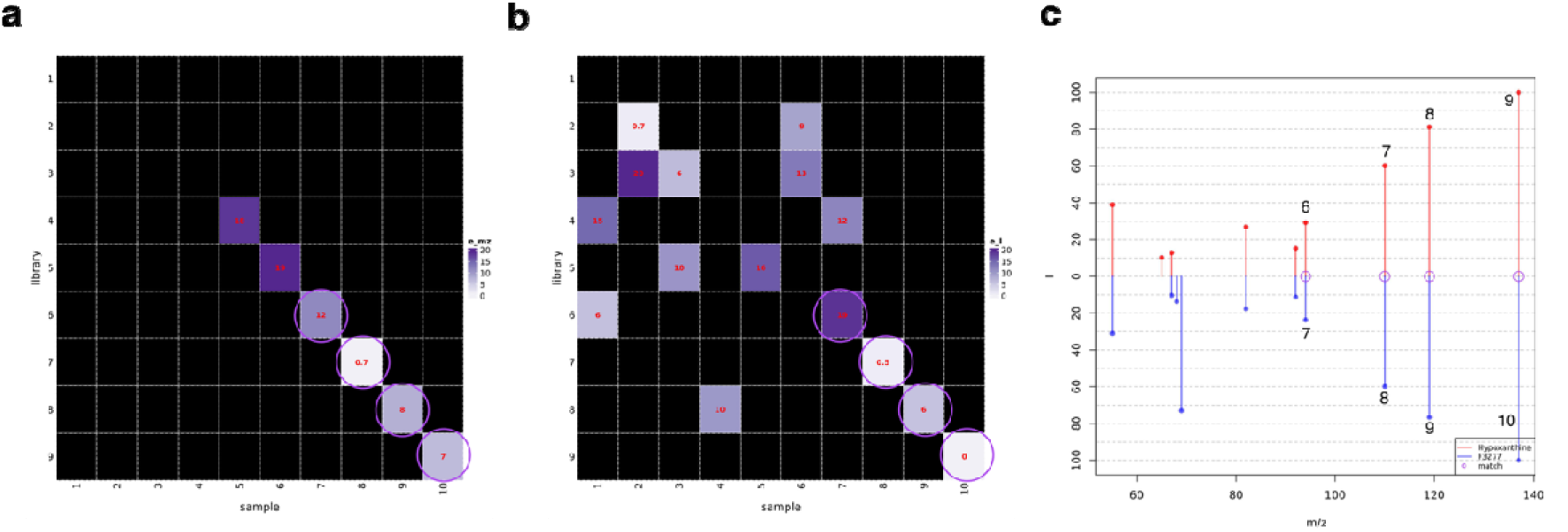
Example plots that show the similarity between observed and reference spectra. **a-b)** ppm error (e_m/z) (a) and relative intensity error (e_I) (b) of all comparisons between peaks in sample (10 MS/MS fragments) and reference library (9 MS/MS fragments) spectra; only the sample-library peak pairs with e_m/z or e_I below 20 are colored in shades of purple, while the other are black; note that in the panel b the sample-library match is only performed based on relative intensity error, regardless of ppm error. **c)** Relative intensity (I) of reference (library) (in red, above zero) and observed (sample) peaks (in blue, below zero) by m/z; numbers close to peaks indicate the fragments of panels a-b. **a-c)** Any pair in which both e_m/z and e_I are below 20 (“match”) is circled in purple.

Lastly, the statistical significance of pathway enrichment for a list of metabolites (represented by PubChem [13] identifiers) can be subjected to over representation analysis and metabolite set enrichment analysis (i.e., gene set enrichment analysis applied to metabolites), by means of a wrapper function that takes advantage of the R package clusterProfiler [14].

## Results

To assess the performance of metabolite identification in margheRita, we set up two experiments (Supplementary Methods, Additional File 1):

i. the first, “Standards”, containing a dataset from a panel of 33 standard metabolites, acquired in IDA mode (information dependent acquisition, which is the DDA mode on SCIEX instrumentation) that could serve as ground truth;
ii. the second, “Urine”, to simulate a “real life dataset” obtained through LC-MS/MS analysis (RP-C18, short gradient, SWATH mode, both polarities) on human urine samples. In both the experiments, we compared the results of margheRita metabolite identification with the annotations provided by MS-DIAL for the same features, knowing that margheRita starts with the advantage of using its own spectral library, while MS-DIAL uses libraries which are much larger in terms of metabolites.

In Standards dataset, margheRita recovered 89% and 82% of the metabolites in, respectively, positive and negative polarity, mostly supported by Level 1 evidence (as expected). We can speculate that margheRita did not reach the 100% recovery because of the possible differences between data extraction, peak picking, peak alignment and peak identification performed by MS-DIAL versus SCIEX-OS [15], which is the SCIEX software used to create the spectral library from true reference standards. When we manually inspected the results, we could verify that few metabolites were not retrieved at level 1 because MS-DIAL algorithm did not assign a good quality MS2 spectrum (i.e. for 12 metabolites in positive polarity, Supplementary Table S5, Additional File 2). Nevertheless, when we checked the corresponding MS2 spectra with SCIEX-OS software, we could univocally assign the identity of those metabolites. In Standards dataset, on the other hand, MS-DIAL correctly annotated the 12% (positive) and 27% (negative) of the corresponding features, mostly at Level 2 (Table 3, Supplementary Tables S5-S6, Additional File 2). In many cases, MS-DIAL left features unknown, without MS2, in both positive and negative polarity (for 29 out of 33 and 24 out of 33 standards, respectively).

**Table 3.**
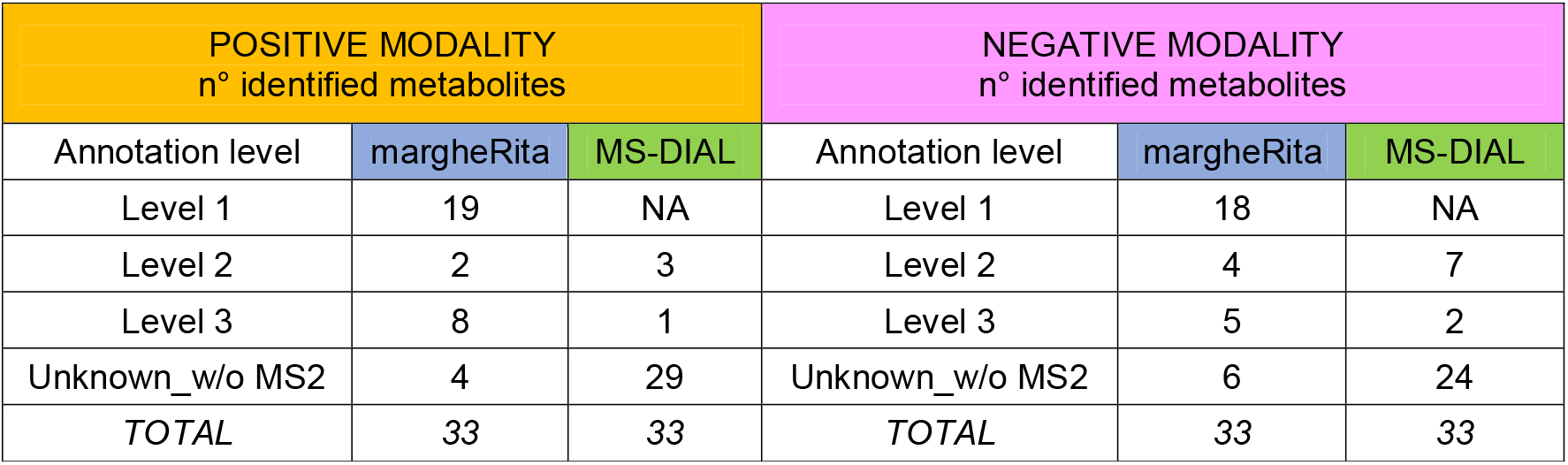
Number of metabolites identified by margheRita in comparison with MS-DIAL v 4.7.0 in “Standards” dataset. Details for level annotation are reported, both for positive (+) and negative (-) polarity using IDA mode (Supplementary Tables S5-S6, Additional File 2). NA: annotation not available; Unknown_w/o MS2: MS-DIAL label for unknown or missing MS2 information.

| POSITIVE MODALITY<br>n° identified metabolites |  |  | NEGATIVE MODALITY<br>n° identified metabolites |  |  |
| --- | --- | --- | --- | --- | --- |
| Annotation level | margheRita | MS-DIAL | Annotation level | margheRita | MS-DIAL |
| Level 1 | 19 | NA | Level 1 | 18 | NA |
| Level 2 | 2 | 3 | Level 2 | 4 | 7 |
| Level 3 | 8 | 1 | Level 3 | 5 | 2 |
| Unknown_w/o MS2 | 4 | 29 | Unknown_w/o MS2 | 6 | 24 |
| *TOTAL* | 33 | 33 | *TOTAL* | 33 | 33 |

In Urine dataset, margheRita identified 386 (of which 135 at Level 1) and 394 metabolites (of which 106 at Level 1) in, respectively, positive and negative polarity, while MS-DIAL provided correct annotations for approximately half of such features (43% and 55% respectively) (Table 4; Supplementary Tables S7-S8, Additional File 2). As expected, these results clearly indicate that the use of an in-house spectral library improves a lot the metabolite annotation, both qualitatively (higher confidence level) and quantitatively (higher number of annotations). Indeed, it is noteworthy that our pipeline is well proficient in assigning MS2 spectra to features which otherwise would remain unknown. Interestingly, we could easily analyze the urine dataset, thus simulating a “real life dataset” obtained by LC-MS/MS SWATH. Indeed, margheRita quickly performed the metabolite annotation step and the statistical analysis to compare the metabolome profiling among the three different donors. As an example, we show (Figure 3) the scaled abundance of the top varying metabolites (ANOVA, FDR q < 0.001) among the three different donors (AA, DD, MM).

**Table 4.** Number of metabolites found by margheRita in comparison with MS-DIAL v 4.7.0 in “Urine” dataset. Details for level annotation are reported, both for positive (+) and negative (-) polarity using SWATH mode Supplementary Tables S7-S8, Additional File 2); NA: annotation not available; Unknown_w/o MS2: MS-DIAL label for unknown or missing MS2 information.

| POSITIVE MODALITY<br>n° identified metabolites |  |  | NEGATIVE MODALITY<br>n° identified metabolites |  |  |
| --- | --- | --- | --- | --- | --- |
| Annotation level | margheRita | MS-DIAL | Annotation level | margheRita | MS-DIAL |
| Level 1 | 136 | NA | Level 1 | 106 | NA |
| Level 2 | 53 | 20 | Level 2 | 16 | 23 |
| Level 3 | 197 | 143 | Level 3 | 272 | 177 |
| Unknown_w/o MS2 |  | 218 | Unknown_w/o MS2 |  | 177 |
| *TOTAL* | *386* | *386* | *TOTAL* | *394* | *394* |

**Figure 3:**
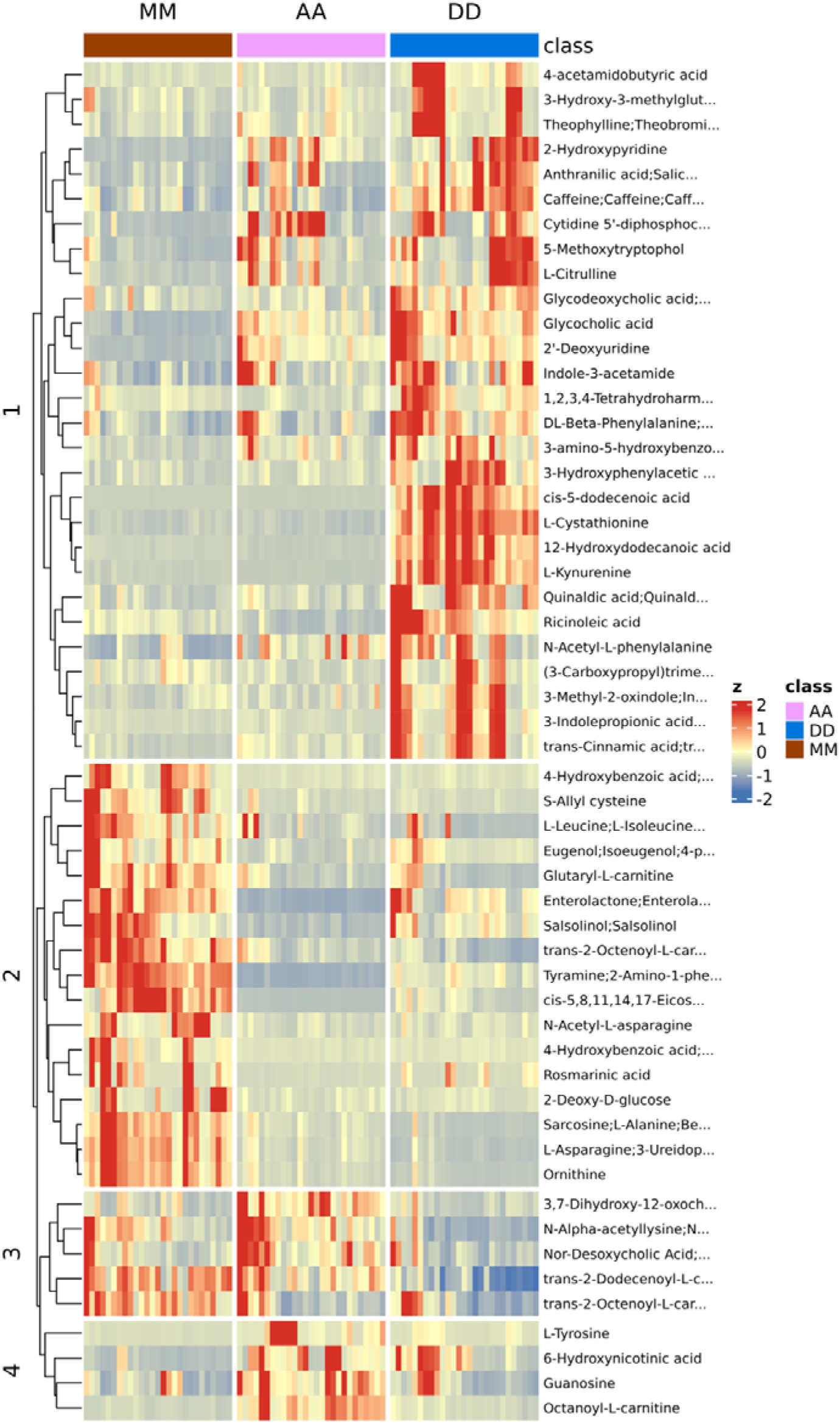
Heatmap of the most variable metabolites among donors in the “Urine” dataset. Scaled abundance of the metabolites that show a significant difference (ANOVA, FDR *q* < 0.001) among the three different donors (AA, DD, MM); metabolite names are trimmed to leave 25 characters; numbers on the left indicate clusters of metabolites with similar pattern of abundance across samples.

## Availability and Future Directions

The R package margheRita (v0.6) is freely available in Zenodo https://doi.org/10.5281/zenodo.20429563, and in GitHub https://github.com/emosca-cnr/margheRita. The accompanying datasets, including the spectral library of authentic chemical standards (rt on the different chromatographic columns, MS and MS/MS data), are available in Zenodo at URL https://doi.org/10.5281/zenodo.17533894.

The development of margheRita addresses a critical need in metabolomics for integrated tools, even capable of handling the complexity of LC–MS/MS SWATH data. Traditional workflows often rely on a combination of heterogeneous software solutions for preprocessing, metabolite identification, and post-acquisition statistical analysis, which can limit reproducibility and increase the risk of analytical bias. By providing a unified, elegant framework that supports both DDA- and DIA-derived datasets, margheRita enhances workflow consistency and facilitates the comparison of results regardless of the acquisition strategy.

While the current version seamlessly integrates with MS-DIAL for initial data processing, our roadmap includes a transition to open MS formats to make this power accessible to an even broader community.

A distinctive strength of the package is the inclusion of an original ‘Level 1’ spectral library of authentic reference standards, which significantly improves the accuracy and confidence of metabolite annotation, a well-known bottleneck in untargeted metabolomics. This resource not only improves the performance of metabolite identification but also contributes to increasing transparency and standardization in metabolomic studies.

Our validation using a SWATH-based dataset demonstrates the robustness and versatility of the package, underlining its potential for widespread application in both exploratory and hypothesis-driven metabolomics research. Beyond its immediate utility, margheRita also provides a foundation for future methodological developments, such as the integration of machine learning approaches for metabolite annotation or the expansion of libraries tailored to specific biological systems. Overall, by streamlining the metabolomics workflow, margheRita contributes to advancing reproducibility, scalability, and interpretability in large-scale metabolomic investigations.

In conclusion, the R package margheRita demonstrates extraordinary versatility; not only does it furnish a comprehensive and multifaceted suite of statistical analyses and sophisticated data representations, but it is also meticulously engineered to be precisely tunable, ensuring it meets most stringent and specific biological requirements.

## Authors’ contributions

EM contributed to project supervision and conceptual guidance, developed the R package, followed the software testing and debugging, performed code optimization and documentation, wrote the original draft and edited the manuscript. MU conceived and designed the study, built the spectral library, performed the experiments. ZA, ENB and VM wrote part of the code, performed code optimization and documentation, contributed to software testing and debugging. GF contributed to code writing, software testing and debugging, and revised the manuscript. DD conceived and designed the study, developed the methodology, performed the experiments, contributed to data interpretation, and edited the manuscript. AA conceived and designed the study, supervised the project, analyzed the data, validated the analytical workflows and verified the results, wrote the original draft, reviewed and edited the manuscript.

