## Supplementary Methods for "MargheRita: streamlining MS-DIAL output analysis and metabolite identification in R"

### **Library**

Sample preparation: The Mass Spectrometry Metabolite Library (MSMLS from SIGMA) contains over 600 unique small molecule metabolites, conveniently provided at 5 µg per well in seven 96 well-plates. The library is intended to be used for mass spectrometry metabolomics applications and provides a broad representation of primary metabolism.

The standards in the plates 1-5 were resuspended in 5% of final volume (up to 20 µL) of high purity methanol (MeOH).

The plates 6 and 7 contained primarily lipid-like compounds (with the exception of the water soluble sugar compounds in plate 6). These compounds were solubilized using a chloroform: methanol (1:1) mixture.

The plate 8 contained bile acids, carnitines, sterols and fatty acids (BACSMLS^TM^, Bile Acid/Carnitine/Sterol Metabolite Library of Standards from SIGMA), providing 5 µg per well. Most compounds were solubilized using high-quality methanol, with the exceptions for the wells: A1, B8, B11, B12, C6, C12 which were solubilized in high-quality chloroform and B10 which was solubilized in high-quality chloroform:methanol (1:1).

The plate 9 contained FAMLS^TM^ (Fatty Acid Metabolite Library of Standards from SIGMA), which is a collection of high quality fatty acids. The standard molecules (all 5 µg per well) in the rows A-C were solubilized in high-quality chloroform, while those ones in the rows D-H were solubilized in high-quality ethanol.

In order to reduce the number of injections, when possible, the 12 standards from each row of the nine 96 well-plates were pooled together.

Data acquisition: Samples were analyzed using the UPLC 1290 (Agilent Technologies) coupled to the TripleTOF 5600+ mass spectrometer (SCIEX) by directly injecting each mixture of standard molecules or single standard molecules.

Chromatographic separations occurred on four different chromatographic columns, using 5 gradients: short and long gradient on RP-C18 column (ACQUITY UPLC HSS T3 Column, Waters, 1.8 µm, 2.1 mm x 100 mm), RP-C8 column (ACQUITY UPLC BEH C8 Column, Waters, 1.7 µm, 2.1 mm x 100 mm), BEH Amide HILIC column (ACQUITY UPLC BEH Amide Column, Waters, 1.7 µm, 2.1 mm x 150 mm), zic-pHILIC column (ZIC®-pHILIC, peek column, MERCK, 5 µm, 2.1 mm x 150 mm).

The short gradient on RP-C18 (RPShort) column used a flow rate set at 0.35 ml/min and a gradient of solvent A (water, 0.1% formic acid) and solvent B (acetonitrile, 0.1% formic acid). The gradient started from 3% B; increased up to 55 % B in 6 min; increased further up to 95 % B in 1.5 min; maintained constant at 95% B for 2 min; decreased to 3% B in 0.5 min and maintained at 3% B for 1 min. The column was set at 45°C while the samples were kept at 4°C. The injection volume was 10 µl. The mass spectrometry analysis was performed both in positive and negative polarity, in the range of m/z 45-750 for the TOF-MS scan with an accumulation time of 150 msec, coupled to an IDA (information dependent acquisition, DDA in SCIEX instrumentation) experiment in the range of m/z 45-750 (the ten most intense ions were selected and fragmented). The mass spectrometer parameters were: Gas 1: 45 psi; Gas2: 50 psi; Curtain gas: 25 psi; Ion Spray Voltage floating (ISVF): 5500 V (-4500 V in negative); Temperature: 500°C; Declustering potential (DP): 80 V (-80V in negative); Collision Energy ± Collision Energy Spread (CE± CES): 35± 15 V (-35 V in negative).

The long gradient on RP-C18 (RPLong) column used a flow rate set at 0.60 ml/min and a gradient of solvent A (water, 0.1% formic acid) and solvent B (acetonitrile, 0.1% formic acid). The gradient started from 2% B; increased up to 95 % B in 14 min; maintained constant at 95% B for 5 min; decreased to 2% B in 0.01 min and maintained at 2% B for 4.99 min. The column was set at 50°C while the samples were kept at 4°C. The injection volume was 10 µl. The mass spectrometry analysis was performed both in positive and negative polarity, in the range of m/z 45-700 for the TOF-MS scan with an accumulation time of 80 msec, coupled to an IDA experiment in the range of m/z 45-700 (the ten most intense ions were selected and fragmented). The mass spectrometer parameters were: Gas 1: 33 psi; Gas2: 58 psi; Curtain gas: 35 psi; Ion Spray Voltage floating (ISVF): 5500 V (-4500 V in negative); Temperature: 500°C; Declustering potential (DP): 80 V (-80V in negative); Collision Energy ± Collision Energy Spread (CE± CES): 40± 15 V (-40 V in negative).

The gradient on RP-C8 column (LipC8) used a flow rate set at 0.40 ml/min and a gradient of solvent A (water, 0.1% formic acid) and solvent B (methanol/isopropanol= 85/15, 0.1% formic acid). The gradient started from 75% B; increased up to 85 % B in 2 min; increased further up to 99.9 % B in 12 min; maintained constant at 99.9% B for 5 min; decreased to 75% B in 1 min and maintained at 75% B for 2 min. The column was set at 50°C while the samples were kept at 4°C. The injection volume was 20 µl. The mass spectrometry analysis was performed both in positive and negative polarity, in the range of m/z 50-1200 for the TOF-MS scan with an accumulation time of 100 msec, coupled to an IDA experiment in the range of m/z 50-1200 (the fifteen most intense ions were selected and fragmented). The mass spectrometer parameters were: Gas 1: 33 psi; Gas2: 58 psi; Curtain gas: 35 psi; Ion Spray Voltage floating (ISVF): 5500 V (-4500 V in negative); Temperature: 500°C; Declustering potential (DP): 80 V (-80V in negative); Collision Energy ± Collision Energy Spread (CE± CES): 45± 15 V (-45 V in negative).

The gradient on BEH Amide HILIC column (HILIC) used a flow rate set at 0.60 ml/min and a gradient of solvent A (acetonitrile, 0.1% formic acid) and solvent B (water, 0.1% formic acid). The gradient started from 2% B; increased up to 60 % B in 10 min; maintained constant at 60% B for 2 min; decreased to 2% B in 0.5 min and maintained at 2% B for 2.5 min. The column was set at 40°C while the samples were kept at 4°C. The injection volume was 15 µl. The mass spectrometry analysis was performed both in positive and negative polarity, in the range of m/z 44-500 for the TOF-MS scan with an accumulation time of 120 msec, coupled to an IDA experiment in the range of m/z 44-500 (the ten most intense ions were selected and fragmented). The mass spectrometer parameters were: Gas 1: 33 psi; Gas2: 58 psi; Curtain gas: 35 psi; Ion Spray Voltage floating (ISVF): 5500 V (-4500 V in negative); Temperature: 500°C; Declustering potential (DP): 80 V (-80V in negative); Collision Energy ± Collision Energy Spread (CE± CES): 35± 15 V (-35 V in negative).

The gradient on zic-pHILIC column (pZIC) used a flow rate set at 0.20 ml/min and a gradient of solvent A (acetonitrile) and solvent B (water, (NH_4_)_2_CO_3_ 20mM, 0.1% NH_4_OH). The gradient started from 20% B; increased up to 80 % B in 15 min; decreased to 20% B in 0.1 min and maintained at 20% B for 5.9 min. The column was set at 45°C while the samples were kept at 4°C. The injection volume was 10 µl. The mass spectrometry analysis was performed both in positive and negative polarity, in the range of m/z 50-1000 for the TOF-MS scan with an accumulation time of 100 msec, coupled to an IDA experiment in the range of m/z 50-1000 (the ten most intense ions were selected and fragmented). The mass spectrometer parameters were: Gas 1: 33 psi; Gas2: 58 psi; Curtain gas: 35 psi; Ion Spray Voltage floating (ISVF): 5500 V (-4500 V in negative); Temperature: 500°C; Declustering potential (DP): 80 V (-80V in negative); Collision Energy ± Collision Energy Spread (CE± CES): 35± 15 V (-35 V in negative).

### **Standards dataset**

The .wiff files, obtained for 33 Standard metabolites, analysed as single metabolites on RP-C18 column with the above reported short gradient and the same parameters for mass spectrometry analysis in positive and negative polarity, were converted in .abf files using the Reifycs Abf Converter and subsequently analysed by MS-DIAL software v. 4.7.0 [1], using parameters reported in Supplementary Tables ST1-2 (Additional File 2). The output file was used to compare the identification performance of “margheRita” and MS-DIAL.

### **Urine dataset**

Untargeted LC-MS/MS metabolomics analysis was performed on different urine samples donated by three healthy participants who voluntarily consumed Kefir shake, Kefir-banana shake and Kefir-papaya shake.

Urine (always midstream), in the first morning and every hour upon kefir shake assumption (N= 8 for each donor for each meal) were collected into a sterile container and placed in an ice bath (2-4°C). The samples were centrifuged immediately at 1,280 g for 30 min at 4 °C to remove any possible cellular contamination. After collection, 10-15 mL of each sample were immediately aliquoted (1 ml each aliquot) and stored at -80°C until analysis. Therefore, in total we collected 72 urine samples, having enrolled three donors fed with three different meals and collected urine every hour for eight hours.

Sample preparation: Samples were prepared normalizing the urine volume. For the preparation of the samples, the urine sample with the smallest volume was considered as reference, so all the rest were diluted with water at the same final volume. To extract the metabolites, 200 µl of urine from each sample were loaded on the Ostro 96 well plate (Waters), filtered, added of 200 µl of acetonitrile filtered again, added of 300 µl of water and filtered. The total collected volume was 700 µl and kept refrigerated at -20°C until further analysis.

Quality controls (QC): Pool of all the extracted urine samples were prepared starting from equal volume of each sample. QC were used to monitor the stability and functionality of the system throughout the instrumental analyses, as they were analysed every ten injections.

Data acquisition: Samples were analyzed using the UPLC 1290 (Agilent Technologies) coupled to the TripleTOF 5600+ mass spectrometer (SCIEX), with the short gradient on RP-C18 column by directly injecting 10 µl of samples. Samples were analyzed in three technical replicates, with randomization. The mass spectrometry analysis was performed both in positive and negative polarity, in the range of m/z 50-850 (TOF-MS scan with an accumulation time of 75 msec), with a SWATH acquisition comprising 16 windows of 50 Da each (accumulation time of 60 msec). The mass spectrometer parameters were: Gas 1: 45 psi; Gas2: 50 psi; Curtain gas: 25 psi; Ion Spray Voltage floating (ISVF): 5500 V (-4500 V in negative); Temperature: 500°C; Declustering potential (DP): 80 V (-80V in negative); Collision Energy ± Collision Energy Spread (CE± CES): 35± 15 V (-35 V in negative).

The obtained .wiff files were converted in .abf files using the Reifycs Abf Converter and subsequently analysed by MS-DIAL software v. 4.7.0 [1], using parameters reported in Supplementary Tables ST3-4 (Additional File 2).

### **Metabolite identification**

The function “metabolite_identification” was run with default parameter values, that is:

| **Description** | **Parameter** | **Value** |
| --- | --- | --- |
| Maximum retention time error | rt_err_thr | 1 |
| Threshold for classifying retention time error as “super | rt_best_thr | 0.5 |
| Threshold for classifying ppm error | accept_flag | 5 |
| Threshold for classifying ppm error | suffer_flag | 10 |
| Threshold for classifying ppm error | unaccept_flag | 20 |
| Minimum relative intensity considered for MS/MS peaks | min_RI | 10 |
| Maximum ppm error for MS/MS peaks | ppm_err | 20 |
| Maximum relative intensity erorr for MS/MS peaks | RI_err | 20 |

### **Filtering of feature-metabolite associations**

When metabolite identification is applied to a dataset of tens of thousands of features from an untargeted metabolomics experiment, it leads to a series of one-to-many and many-to-one associations between features and metabolites. The following algorithm leverages the four error types and the feature-metabolite association classification to disentangle such relations and identify the associations supported by the strongest evidence. The algorithm is composed of four steps:

1. To resolve the one-to-many associations between ${(f}_{i},m_{j})$ pairs and MS/MS spectra $s_{jh}$, for each pair ${(f}_{i},m_{j})$, select the spectrum $s_{jh}$ that has:
   a. the highest number $n_{ij}$ of matches between feature MS/MS fragments and metabolite MS/MS fragments (“peaks_found_ppm_RI” column);
   b. the highest ratio $n_{ij}/n_{l}$ where $n_{l}$ is the number of metabolite MS/MS fragments (“matched_peaks_ratio” column).
2. To resolve the one-to-many associations between $m_{j}$ and $f_{i}$, for each $m_{j}$, select the $f_{i}$ that has:
   a. the highest $n_{ij}/n_{l}$ ;
   b. the best annotation level (“Level” column);
   c. the lowest $\varepsilon_{m/z}\left( f_{i},m_{j} \right)$ (“ppm error” column).
3. To resolve the one-to-many associations between $m_{j}$ and $f_{i}$ not supported by MS/MS spectra, for each $m_{j}$, select the $f_{i}$ that has:
   a. the best annotation level (“Level” and “Level_note” columns);
   b. the best mass match (“mass_status” column);
   c. the best retention time match (“RT_class” column);
   d. the lowest $\varepsilon_{\mathrm{rt}}\left( f_{i},m_{j} \right)$ (“RT_err” column);
   e. the lowest $\varepsilon_{m/z}\left( f_{i},m_{j} \right)$.
4. To resolve the one-to-many associations between $f_{i}$ and $m_{j}$, for each $f_{i}$ keep the $m_{j}$ that has:
   a. the best annotation level;
   b. the best mass match;
   c. the highest $n_{k}^{*}$;
   d. the best retention time match;
   e. the highest $n_{k}^{*}/n_{l}$;
   f. the lowest $\varepsilon_{\mathrm{rt}}\left( f_{i},m_{j} \right);$
   g. the lowest $\varepsilon_{m/z}\left( f_{i},m_{j} \right)$.
